# Royal knifefish generate powerful suction feeding through large neurocranial elevation and high epaxial muscle power

**DOI:** 10.1101/2022.01.17.476172

**Authors:** Ellen Y. Li, Elska B. Kaczmarek, Aaron M. Olsen, Elizabeth L. Brainerd, Ariel L. Camp

## Abstract

Suction feeding in ray-finned fishes involves powerful buccal cavity expansion to accelerate water and food into the mouth. Previous XROMM studies in largemouth bass (*Micropterus salmoides*), bluegill sunfish (*Lepomis macrochirus*), and channel catfish (*Ictalurus punctatus*) have shown that more than 90% of suction power in high performance strikes comes from the axial musculature. Thus, the shape of the axial muscles and skeleton may impact suction feeding mechanics. Royal knifefish (*Chitala blanci*) have an unusual postcranial morphology, with a ventrally flexed vertebral column and relatively large mass of epaxial muscle. Based on their body shape, we hypothesized that royal knifefish would generate high power strikes by utilizing large neurocranial elevation, vertebral column extension, and epaxial shortening. As predicted, *C. blanci* generated high suction expansion power compared to the other three species studied to date (up to 160 W), which was achieved by increasing both the rate of volume change and the intraoral subambient pressure. The large epaxial muscle (25% of body mass) shortened at high velocities to produce large neurocranial elevation and vertebral extension (up to 41 deg, combined), as well as high muscle mass-specific power (up to 800 W kg^-1^). For the highest power strikes, axial muscles generated 95% of the power, and 64% of the axial muscle mass consisted of the epaxial muscles. The epaxial-dominated suction expansion of royal knifefish supports our hypothesis that postcranial morphology may be a strong predictor of suction feeding biomechanics.

**SUMMARY STATEMENT:** Royal knifefish rely on their distinct postcranial morphology--with a curved vertebral column and large dorsal body muscles--to produce large neurocranial elevation and powerful suction feeding.

## INTRODUCTION

High performance suction feeding in ray-finned fishes is both fast and forceful, requiring high power to expand the buccal cavity and suck in prey. Instantaneous suction expansion power can be measured empirically by using X-ray Reconstruction of Moving Morphology (XROMM) to measure instantaneous buccal cavity volume and rate of buccal cavity expansion (Camp et al., 2015). Combined with measurements of subambient buccal pressure, buccal volume measurements make it possible to calculate instantaneous suction expansion power as the product of rate of buccal volume change and subambient buccal pressure (Van Wassenbergh et al., 2015).

To date, suction expansion power has been measured with XROMM in three species of ray-finned fishes: largemouth bass (*Micropterus salmoides*), bluegill sunfish (*Lepomis macrochirus*), and channel catfish (*Ictalurus punctatus*) (Camp et al., 2015; Camp et al., 2018; Camp et al. 2020). In the highest performance strikes from all three species, the empirically measured suction power was far too great to have been generated by muscles in the head region alone. Instead, more than 90% of suction power came from epaxial and hypaxial musculature (largemouth bass and bluegill sunfish) or the hypaxial musculature (channel catfish). Furthermore, the axial musculature was found to actively shorten along 60-70% of the length of the body, encompassing the majority of axial muscle mass (Camp et al., 2015; Camp et al., 2018; Camp et al., 2020; Jimenez and Brainerd, 2020; Jimenez and Brainerd, 2021). Thus, although it has long been known that axial musculature contributes to suction feeding (Liem, 1967; Osse, 1969), the ability to measure suction power, muscle length, and activation has revealed that some fish use nearly their whole bodies for suction feeding. The overall body shape and musculoskeletal morphology should therefore be considered when studying the biomechanics and energetics of suction feeding (Camp and Brainerd, 2022).

There are several ways the morphology of the body and axial muscles, including skeletal elements linking the head and body, can impact intraoral pressure, buccal volume, and ultimately suction power. First, the shape of the body reflects the relative size and distribution of the axial muscles, which may impact their function during feeding. Carroll et al. (2004) found that deeper-bodied fish had greater epaxial cross-sectional area and longer epaxial moment arms for cranial elevation. As a result, deep-bodied bluegill sunfish were capable of greater pressure generation during feeding than the more fusiform largemouth bass (Carroll et al., 2004). Both the dorsoventral depth (Fig. 1C) and transverse shape (Fig. 1D) of the body reflect the relative cross-sectional area of the epaxial and hypaxial muscles. While hypaxial muscles typically have smaller cross-sectional areas anteriorly where they surround the body cavity, these muscles contribute substantially to suction power in all species studied so far with XROMM.

**Fig. 1.**
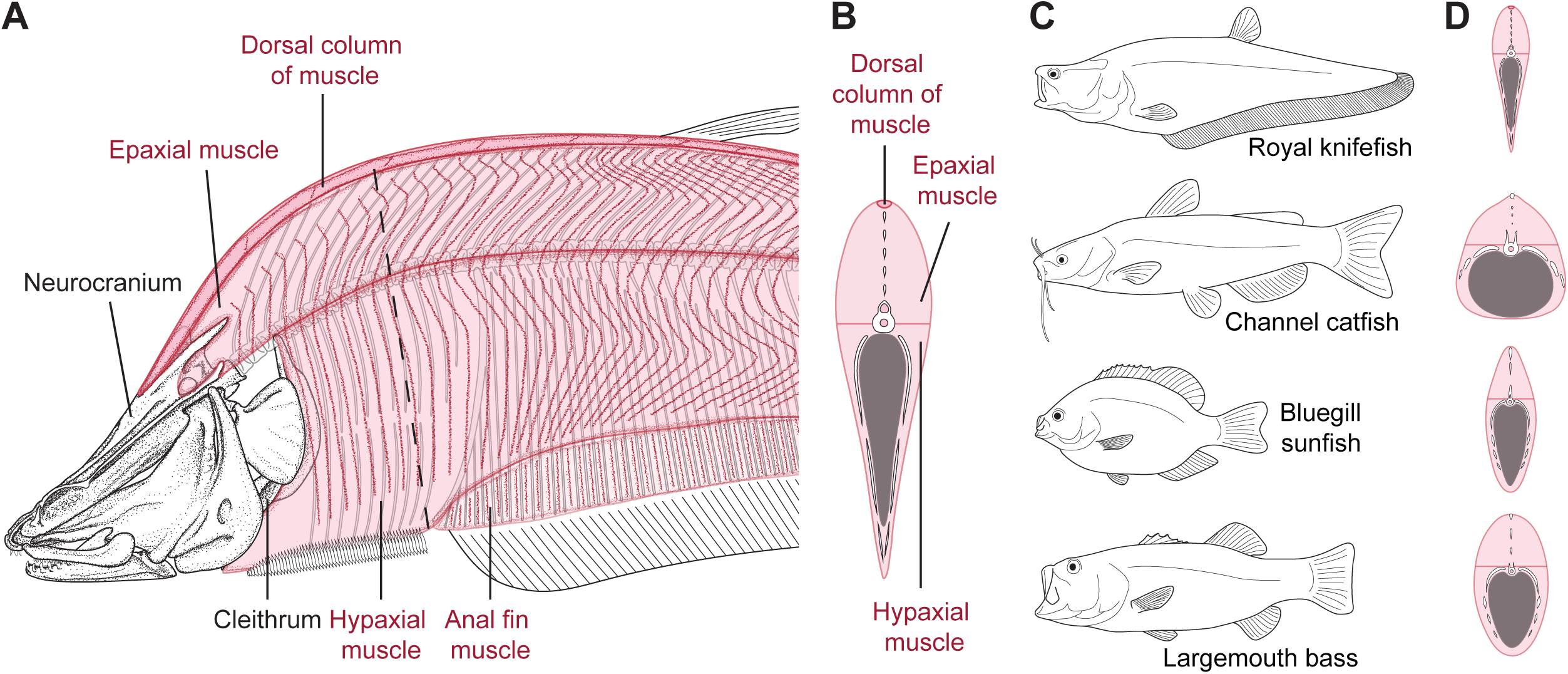
Body shape and anatomy of the axial musculature and skeleton in royal knifefish, *Chitala blanci*, compared with the body shapes of three other species for which suction power has been measured. (A) Lateral view of the neurocranium, cleithrum, and left-side bones of the head in *C. blanci*. Dashed line indicates the approximate location of the cross-section shown in (B). (B) Transverse cross-section of *C. blanci*, illustrating its inverted teardrop shape. (C) Whole-body shape and (D) transverse cross-section comparisons of *C. blanci* to other previously studied species: channel catfish (*Ictalurus punctatus*), bluegill sunfish (*Lepomis macrochirus*), and largemouth bass (*Micropterus salmoides*). Fishes are shown with expanded mouths in (C).

Second, body shape and skeletal anatomy may influence neurocranial elevation, a common component of mouth expansion and an essential motion for transmitting epaxial muscle power to the head. Deep-bodied fish with more bony processes (i.e. supraneurals, neural spines, and pterygiophores) immediately caudal to the neurocranium—like bluegill sunfish—had less neurocranial elevation than largemouth bass (Jimenez et al., 2018; Camp and Brainerd, 2014; Camp et al., 2018; Table 2). The channel catfish, which has even more postcranial ossifications, uses little or no neurocranial elevation (Camp et al., 2020). Rather, catfish relied on hypaxial muscle power, transmitted via retraction of their robust pectoral girdle (Camp et al., 2020). These inter-species comparisons demonstrate emerging links between postcranial morphology and suction feeding power and biomechanics. However, so far only a small sample of body shapes and species have been investigated.

The royal knifefish (*Chitala blanci*) offers an interesting model for studying suction feeding, as it is both morphologically and phylogenetically distinct from species previously studied with XROMM. The royal knifefish is a member of the family Notopteridae (Order Osteoglossiformes) and is not closely related to channel catfish (Order Siluriformes) nor largemouth bass and bluegill sunfish (Order Centrarchiformes). Morphologically, royal knifefish have a ventrally flexed vertebral column and depressed neurocranium in their resting posture, dorsoventrally deep epaxial musculature, and laterally compressed body (Fig. 1; Coombs and Popper, 1982; Sanford and Lauder, 1989). In addition, royal knifefish have an inverted teardrop shaped transverse cross-section, with the body being thickest at the epaxial muscles and tapering in thickness ventrally towards the hypaxials and anal fin (Fig. 1B).

Of the species previously studied with XROMM, bluegill sunfish have the most similar body shape to royal knifefish. Both species have laterally compressed and deep bodies, ventrally curved vertebral columns, and dorsoventrally deep epaxial muscles (Fig. 1). Similar to bluegill sunfish, the epaxial muscles of royal knifefish provide a relatively large cross-sectional area and large moment arm, which may enable them to generate similarly large subambient buccal pressures. Compared to largemouth bass and channel catfish, bluegill sunfish generated the most powerful suction expansion relative to their body and muscle mass, by generating greater subambient buccal pressures with smaller axial muscles (Camp et al., 2018). Since royal knifefish have similar epaxial morphology, we expect they can also generate powerful suction expansion, relative to their muscle mass.

Royal knifefish also differ from bluegill sunfish in key ways, which we hypothesize will result in greater neurocranial elevation and epaxial contribution in royal knifefish. Royal knifefish have a more laterally compressed and craniocaudally elongated head and body, a more ventrally flexed vertebral column, fewer bony processes caudal to the neurocranium, and a greater proportion of epaxial muscle than bluegill sunfish. The exaggerated ventral flexion of the vertebral column causes the neurocranium to have a depressed resting posture, which we expect increases its range of dorsoventral motion (Fig. 1). Additionally, the curvature may cause the axis of rotation of the neurocranium to be located more caudally (close to the vertebral column inflection point), which has also been correlated with greater neurocranial elevation (Jimenez et al., 2018). Compared to bluegill sunfish, royal knifefish have few bones immediately caudal to the neurocranium: no supraneurals or dorsal fin pterygiophores, and thin neural spines. We predict that this enables them to perform larger neurocranial elevation than bluegill sunfish. Lastly, while both royal knifefish and bluegill sunfish have dorsoventrally deep epaxial muscles, the transverse cross-section of the royal knifefish (forming an inverted teardrop) increases the epaxial muscle mass relative to the hypaxials (Fig. 1). Based on the body shape of the royal knifefish, we hypothesize that they rely predominantly on massive epaxial muscles and large neurocranial elevation to power suction feeding.

To test these hypotheses, we used XROMM to measure the 3D skeletal kinematics and instantaneous buccal volume of royal knifefish during suction feeding. Intraoral pressure was also measured simultaneously and combined with the rate of buccal volume change to calculate the suction power during royal knifefish strikes (Camp et al., 2015; Camp et al., 2020). Length changes were measured throughout the epaxial, hypaxial, and sternohyoid muscles during suction feeding using fluoromicrometry (Camp et al., 2016). Muscle shortening and post-mortem muscle mass were used to determine the roles and relative contributions of these muscles to suction power. These data allowed us to test if royal knifefish 1) have relatively large cranial elevation compared to previously studied species (bass, sunfish, and catfish) and 2) predominantly utilize epaxial muscle power when suction feeding. Determining how royal knifefish use their unusual postcranial morphology to power suction expansion provides a better understanding of the relationship between body shape and suction feeding biomechanics.

## MATERIALS AND METHODS

Royal knifefish (*Chitala blanci*, d’Aubenton 1965) were acquired from Ocean State Aquatics, Coventry RI: Cb01 (standard length 35.6 cm, body mass 217 g), Cb03 (30.8 cm, 170 g), and Cb04 (43.3 cm, 480 g). Royal knifefish were maintained on a diet of goldfish (*Carassius auratus*). All experimental procedures were approved by Brown University Institutional Animal Care and Use Committee.

The fish were anesthetized with a buffered MS-222 solution during surgical implantation of a buccal cannula for pressure measurement and radio-opaque bone and muscle markers. Implantation techniques were consistent with those previously reported (Camp and Brainerd, 2014), and are described here in brief. One to five radio-opaque markers (tantalum spheres 0.50 or 0.80 mm in diameter) were implanted into the neurocranium, the left and right ceratohyals and cleithra, and the left maxilla, lower jaw, suspensorium, and operculum (Fig. 2A,B). Cb04 received bilateral lower jaw implantations. In all individuals, 0.80 mm tantalum beads were implanted superficially, slightly to the left of the mid-sagittal plane in the epaxial (five to nine markers), and sternohyoid musculature (two to three markers) (Fig. S1). Ventral muscles were marked in Cb01 (anal fin) and Cb04 (hypaxial) with three to five markers, with no ventral markers in Cb03. The dorsal column of epaxial musculature was implanted in Cb04 (three markers) (Fig. S1). Five to six muscle markers were used to define a body plane (Fig. S1). Following established methods, a cannula guide for the pressure transducer was implanted into the ethmo-frontal region of the neurocranium, avoiding the palatine and teeth, protruding just into the buccal cavity (Norton and Brainerd, 1993). All individuals received perioperative analgesic (butorphanol or ketoprofen) and Cb01 and Cb03 received an antibiotic (enrofloxacin). Fish were allowed to recover fully, i.e., resumed natural and aggressive feeding behaviors, before filming experiments began.

**Fig. 2.**
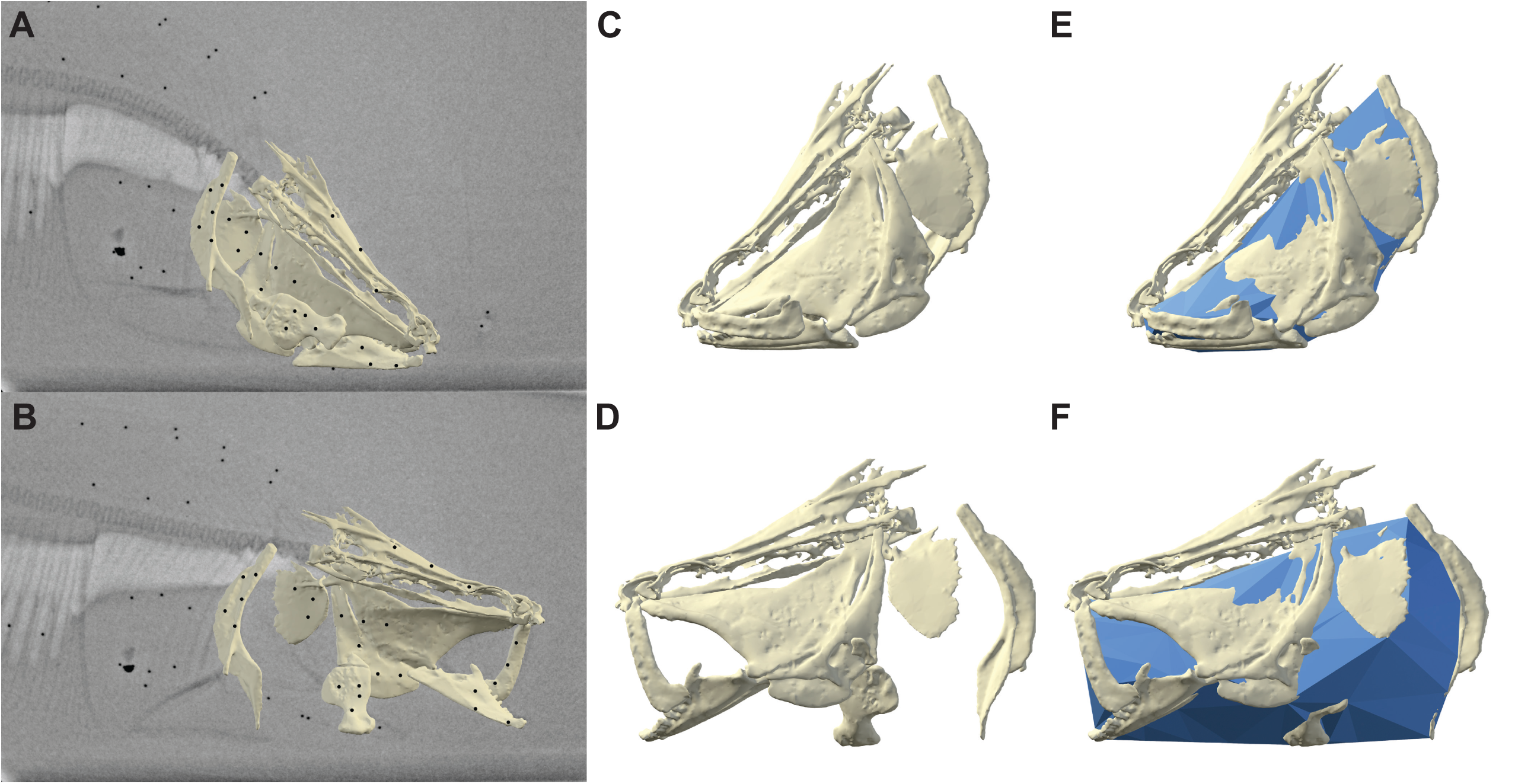
Sample XROMM animation and measurement of buccal cavity volume before and during suction expansion. Animated bone models before suction expansion (A,C,E) and during suction expansion (B,D,F). (A,B) Right lateral view of X-ray image with animated neurocranium, left cleithrum, and left-side bone meshes. Surgically implanted bone and intramuscular (epaxial, hypaxial, dorsal column, and sternohyoid) markers are visible as black circles. (C,D) Left lateral view of animated bone models and (E,F) animated dynamic endocast of buccal cavity volume.

### Data recording

All fish were trained to feed on live goldfish (approximately 3-5 cm total length) in custom-built acrylic aquaria with a feeding extension tunnel (75–100 mm wide, 300-400 mm long) designed to minimize the amount of water through which the X-ray beams must travel (Gidmark et al., 2012). See Movie S1 for a standard light video (recorded at 500 frames s^-1^ and slowed down 16.7 times) of Cb04 feeding in a tunnel.

A custom biplanar X-ray system (Imaging Systems and Services, Painesville, OH, USA) was used to capture dorsoventral and lateral X-ray videos at 500 frames s^-1^ with Phantom v10 high-speed cameras (Vision Research, Wayne, NJ, USA) at 100 mA and 90-115 kV. Standard grid and calibration objects were used to remove distortion introduced by X-ray machines and to calibrate three-dimensional (3D) space (Brainerd et al. 2010). Intraoral pressure was simultaneously recorded with an SPR-407 Mikro-tip pressure probe (Millar Instruments, Houston, TX, USA) inserted into the neurocranial cannula, recording at 1000 Hz with PowerLab and LabChart 7.2.2 (ADInstruments, Colorado Springs, CO, USA). Pressure transducer calibration was carried out daily by moving the probe through a 10 cm change in water depth while recording the voltage output. This model of probe provides linear pressure-voltage outputs over a pressure range of at least 0 to -60 kPa (Higham et al., 2006). Pressure data were collected for each strike and noise was filtered in R (2019, R Core Team, Vienna, Austria) using a low-pass, forward-backward (to remove phase shifts) Butterworth filter at a cutoff frequency of 200 Hz. The pressure recording was initiated by the X-ray camera trigger and corrected for a mean measured lag of two milliseconds (range 1-3 ms) from the initial X-ray image. For three Cb03 strikes, the initial X-ray images were missing, so to correctly align the pressure and video data, we averaged the time between peak pressure and peak rate of volume change for all Cb04 strikes and corrected the Cb03 image sequences to account for a pressure lag of two milliseconds. Feeding trials with the greatest subambient buccal pressure from each individual were chosen for analysis. A total of 23 recorded strikes (six from Cb01, seven from Cb03, ten from Cb04) were analyzed.

Computed tomography (CT) scans were taken of each fish after surgical implantation with a FIDEX CT Scanner (Animage, Pleasanton, CA, USA), with 480 × 480 pixel resolution and 0.173 mm slice thickness. From these scans, polygonal meshes of each bone and the radio-opaque markers were generated in Horos (v3.3.5; Horos Project; horosproject.org) and edited in Geomagic 2014 (Research Triangle Park, NC, USA). Markers were imported into Autodesk Maya 2020 (San Rafael, CA, USA) and with custom scripts from ‘XROMM Maya Tools’ package--available at https://bitbucket.org/xromm/xromm_mayatools--their respective *xyz* (3D) coordinates were determined. Raw data for this study are publicly available and stored on the XMAPortal (http://xmaportal.org) in the study “Knifefish Suction Feeding,” with the permanent identifier BROWN65. Video data are stored with their essential metadata in accordance with best practices for video data management in organismal biology (Brainerd et al., 2017).

### XROMM animation

For each of the three individual *C. blanci*, skeletal kinematics were reconstructed using marker-based XROMM with XMALab 1.5.5 (Knörlein et al., 2016; software and instructions available at https://bitbucket.org/xromm/xmalab) and custom XROMM MayaTools scripts. Markers from both X-ray videos were tracked in XMALab with a mean precision of 0.05 mm and maximum precision error of 0.1 mm across all trials (measured as the standard deviation of the unfiltered pairwise marker-to-marker distances within all rigid bodies). Marker coordinates were used to reconstruct the 3D motion of each bone using the ‘matools’ R package, following the XROMM workflow described in Olsen et al., 2019 (available under matools R package at https://github.com/aaronolsen). Briefly, all *xyz* marker coordinates were smoothed and, for bones containing three or more markers, 3D coordinates were combined with their respective CT coordinates (using the ‘unifyMotion’ function from ‘matools’) to produce rigid body transformations. These transformations were applied to the skeletal bone meshes in Maya (2020, Autodesk), producing a 3D XROMM animation of each suction feeding strike (Fig. 2A,B). For any bones with only two beads or those with a linear set of markers, virtual constraints were applied using the ‘matools’ R package in accordance with anatomical constraints (e.g. cartilaginous symphysis between the cranioventral region of the left and right cleithra, or ceratohyal retraction along the neurocranial mid-sagittal plane).

The body plane was animated with a set of five to six intramuscular axial markers in roughly the same location along the body for each individual. These markers were positioned near the most curved region of the vertebral column (Fig. S1). Their 3D coordinates were combined with their respective CT coordinates to generate a rigid body transformation using the ‘matools’ R package (Olsen et al., 2019). The body plane animation was included in the XROMM animations mentioned above.

### Skeletal kinematics

Six-degree-of-freedom motions of the neurocranium and left cleithrum were measured relative to the body plane. These rotations were measured with a joint coordinate system (JCS), which measures the relative rotations of two anatomical coordinate systems (ACSs), one attached to the bone and the other to the body plane (Camp and Brainerd, 2014; Camp et al., 2018). Each JCS measured translation and Euler angle rotations about the x-, y-, z-axis, following the right-hand rule and *zyx* order of rotation. The JCS used to measure neurocranial motion was placed at the craniovertebral joint and the JCS to measure cleithral motion was placed at the dorsal tip of the cleithra. Both sets of JCSs were aligned with the z-axis oriented mediolaterally, y-axis rostrocaudally, and the x-axis dorsoventrally. Z-axis rotations were standardized to start at 0 deg by subtracting their value at the start of each strike. Positive rotation about the z-axis reflects dorsal rotation in the sagittal plane, corresponding to neurocranial elevation or cleithral protraction. Rotations about the z-axis also reflect dorsoventral motions of the cranial vertebrae, as these impact the position and motion of the body plane.

### Dynamic endocast

Following previously established methods, changes in buccal cavity volume were measured from XROMM animations using a dynamic endocast (Camp et al., 2015; Camp et al., 2020). In brief, a polygonal mesh endocast of the left side of the buccal cavity was generated using locators attached to the inside surface of cranial bones. Additional locators were placed between bones to define the ventral border of the buccal cavity, i.e., the sternohyoid and protractor hyoideus muscles, and the mid-sagittal plane dividing the left and right sides of the buccal cavity. The 3D coordinates of the locators were imported into MATLAB (R2020a; MathWorks, Natick, MA, USA) and custom-written scripts (available at https://bitbucket.org/ArielCamp/dynamicendocast) were used to generate the volume enclosing the locators and calculate its volume, for each frame. For each frame, the volume was generated from the xyz coordinates of the locators using an alpha shape: a method of fitting or “wrapping” 3D points with a 3D shape (Edelsbrunner et al., 1983). Alpha shapes are a generalization of convex hulls that allow the fineness of fit to be varied by changing the alpha value and allows the shape to include concave curvatures. The volumes were generated using the ‘alphashape’ function in MATLAB, and an alpha value of 3 was found to provide the best fit, i.e., endocasts fully filled the mouth cavity with minimal interpenetration of the bone models. Polygonal meshes (.obj files) of the volumes of the left side of the buccal cavity were then imported into Maya for visual verification (Fig. 2E,F). Under assumptions of bilateral cranial symmetry, the left mesh volume was doubled to calculate bilateral buccal volume expansion.

### Muscle length changes

Sternohyoid and axial muscle length changes were measured from X-ray videos as the distance between intramuscular markers, i.e., by fluoromicrometry (Camp et al., 2016). Muscle markers were tracked in XMALab and their coordinates were filtered in R with ‘matools’ as described above. The distance between muscle markers was subsequently calculated in R to determine the magnitude and distribution of sternohyoid, epaxial, and hypaxial muscle shortening. To capture muscle shortening in the cranialmost region of the epaxial muscle and the dorsal column of Cb04, the distance was measured between the first marker in the muscle region and a locator placed at the cranialmost neurocranium-epaxial muscle attachment site of the animated neurocranium model in Maya. Distance in the cranialmost hypaxial region was calculated between the first hypaxial muscle marker and a locator attached to the cleithra, placed in line with the hypaxial muscle bead set (Fig. S1). Since the entire marker set was not consistently within the X-ray imaging volume, axial muscle length was measured from a subset of axial muscle markers that were visible in almost all strikes. This set of markers extended approximately 7-9 cm caudal of the craniovertebral joint: from the neurocranium to as far back as a few centimeters cranial of the dorsal fin (Fig. S1). Within this region, fluoromicrometry was used to estimate the muscle lengths of subregions along the length of the body by measuring distance between adjacent pairs of intramuscular markers.

For the axial muscles, whole-muscle length was calculated by taking the sum of the subregional muscle lengths, originating with the cranialmost locator and extending to the caudalmost visible muscle marker. In the sternohyoid, all implanted markers were visible, and its measurements are reported as whole-muscle length. Muscle length at each time step was normalized by the mean initial length measured at the first recorded frame of each strike (*L*_i_), with values less than one representing that the muscle had shortened. Muscle velocity was similarly calculated at each time as the change in normalized muscle length divided by the change in time, denoted by *L*_i_ s^-1^, with positive values representing muscle shortening. Note that this method for determining axial muscle strain differs slightly from other suction power studies, in that we used the sum of the subregional muscle lengths, whereas prior papers used the distance from the neurocranium or cleithrum to the caudalmost axial muscle marker (Camp et al., 2015; Camp et al., 2018; Camp et al., 2020). The summation method recorded more consistent levels of epaxial muscle shortening in *C. blanci*, likely due to its ability to capture the length of the naturally flexed epaxial musculature at rest.

### Power calculations

Instantaneous suction power was calculated in R as the product of rate of volume change and intraoral pressure as described in Camp et al. (2015). Before calculating rate of volume change, bilateral buccal volume measurements from the dynamic endocast were filtered with a low-pass, forward-backward Butterworth filter (150 Hz cutoff) to reduce noise generated by frame-to-frame polygonal mesh re-triangulations. Buccal pressure was downsampled from 1000 Hz to 500 Hz to match the frequency of the volume data. Pressure data were calculated relative to initial, ambient pressure prior to the strike and multiplied by -1, so that at each time step, the product of subambient pressure and increasing rates of volume change would reflect positive power (Fig. 3).

**Fig. 3.**
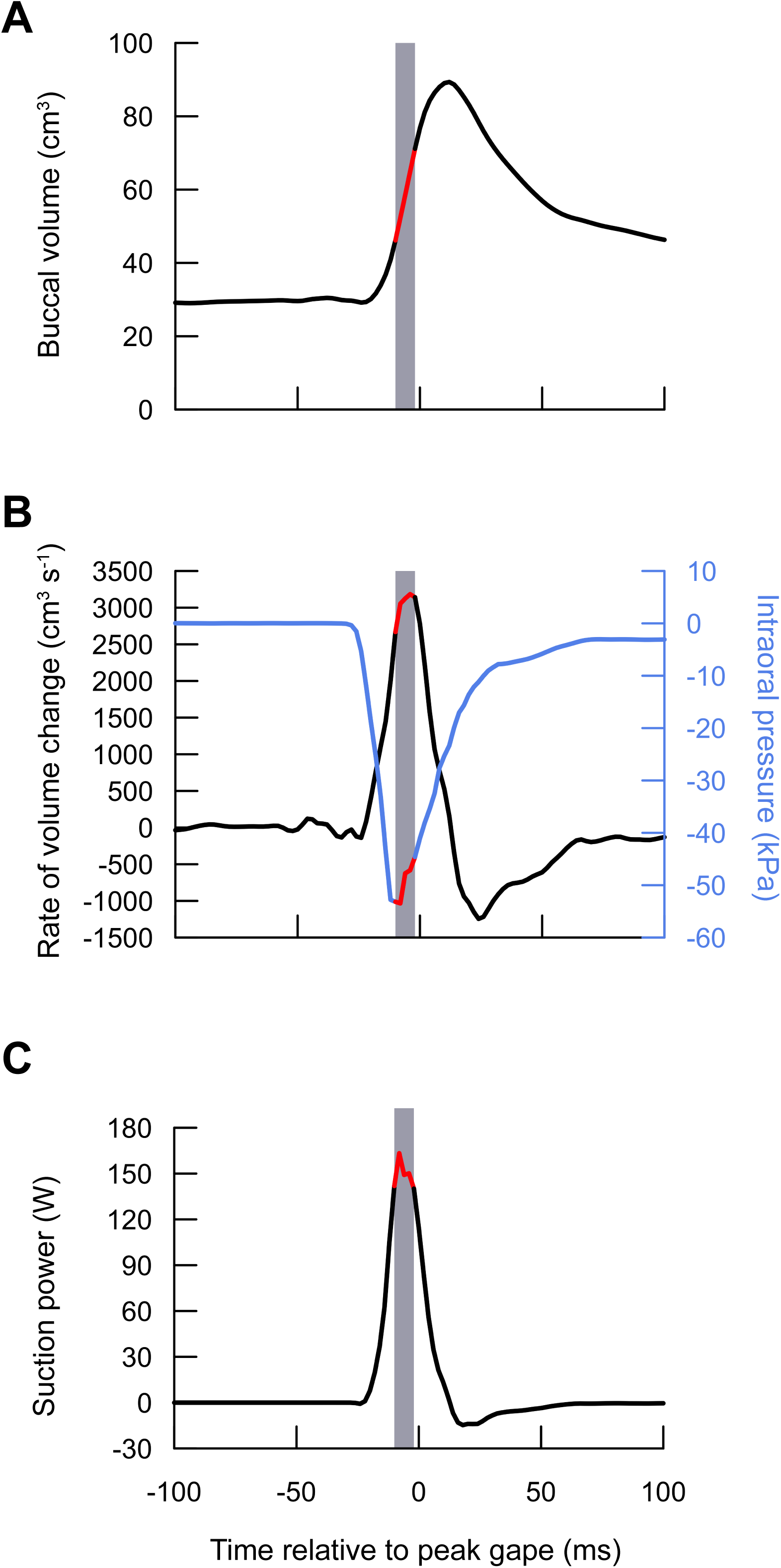
Buccal volume, rate of volume change, intraoral pressure, and suction power in the highest power strike. (A) Buccal volume, measured using dynamic endocast. (B) Rate of volume change (black, left axis) and intraoral pressure relative to ambient pressure (blue, right axis). (C) Suction power was calculated as the product of intraoral pressure and rate of volume change at each time point. Peak suction power (within 25% of maximum) is indicated by the shaded region with the corresponding values highlighted in red.

For each strike, axial and cranial mass-specific power were calculated by dividing the maximum instantaneous power by the mass of the respective muscle groups. Muscle masses were determined by post-mortem dissection of the muscle regions on the right side of the fish, weighed on a digital scale, and then doubled to estimate bilateral muscle mass for all individuals except Cb03. The body of Cb03 was unavailable for dissection, so muscle masses are estimates determined by averaging the percent of muscle mass for each muscle of Cb01, Cb04, and an additional individual, Cb02, and assuming proportionality based on overall body mass across individuals (Table 1). The total body mass, and bilateral epaxial, hypaxial, dorsal column, sternohyoid, and cranial muscle masses from Cb02 were 0.393 kg, 0.103 kg, 0.066 kg, 0.002 kg, 0.0032 kg, and 0.0054 kg respectively (values for other individuals are reported in Table 1). In accordance with previous XROMM studies, epaxial muscle mass included all of the epaxial musculature dorsal to the vertebral column and about 60-70% along the length of the body, based on the extent of shortening identified (Camp and Brainerd, 2014; Camp et al., 2018; Camp et al., 2020; Jimenez and Brainerd, 2020; Jimenez et al., 2021). Note that this method differs from prior studies, which only included epaxial musculature dorsal to the cleithrum-supracleithrum joint, and which reported lower percentages of epaxial muscle mass in bluegill sunfish and largemouth bass (Carroll, 2004; Carroll and Wainwright, 2009). Axial mass-specific power was calculated by dividing instantaneous power by the sum of the epaxial, hypaxial, and dorsal column muscle mass. Cranial mass-specific power was calculated by dividing instantaneous power by the combined mass of the levator arcus palatini, dilator operculi, levator operculi, and sternohyoid muscles. Muscle mass-specific power was determined by dividing the maximum instantaneous power by the total mass of the muscle regions (epaxial, hypaxial, and sternohyoid muscle) shortening during suction expansion (Fig. 3). These mass-specific values represent the estimated amount of power each group of muscles would need to output if they were the sole contributors to suction feeding expansion.

**Table 1.** Mean (± s.e.m) measurements of peak pressure, change in buccal volume, power, total body mass, bilateral axial and sternohyoid muscle mass, and peak muscle mass-specific power of each individual. Mean peak axial and sternohyoid muscle strain and shortening velocity were measured during the period of peak power.

| Variable | Cb01 (N = 6) | Cb03 (N = 6) | Cb04 (N = 10) |
| --- | --- | --- | --- |
| Peak pressure, kPa | $-7.9 \pm 2.0$ | * $-15.2 \pm 4.5$ | $-44.2 \pm 3.5$ |
| Peak change in buccal volume, $\text{cm}^3$ | $23.4 \pm 3.5$ | * $15.9 \pm 2.5$ | $54.2 \pm 3.4$ |
| Peak rate of volume change, $\text{cm}^3 \text{s}^{-1}$ | $1000 \pm 239$ | * $829 \pm 178$ | $2518 \pm 127$ |
| Peak power, W | $6.5 \pm 2.1$ | * $15.9 \pm 6.2$ | $100.6 \pm 10.5$ |
| Total body mass, g | 217 | 170 | 480 |
| Epaxial |  |  |  |
| ▲ Muscle mass, g | 52.7 | ** 44.1 | 125.2 |
| Initial muscle length, mm | $115.1 \pm 0.6$ | $109.4 \pm 0.5$ | $103.4 \pm 0.2$ |
| Peak muscle strain, % | $3.4 \pm 0.4$ | $7.9 \pm 1.4$ | $8.0 \pm 0.7$ |
| Peak muscle velocity, $\text{L}_i \text{s}^{-1}$ | $2.0 \pm 0.2$ | $4.6 \pm 1.5$ | $4.7 \pm 0.1$ |
| Hypaxial |  |  |  |
| Muscle mass, g | 27.5 | ** 25.5 | 73.7 |
| Initial muscle length, mm | *** $100.4 \pm 3.0$ | **** | $65.3 \pm 0.5$ |
| Peak muscle strain, % | *** $1.8 \pm 2.8$ | **** | $-1.1 \pm 1.3$ |
| Peak muscle velocity, $\text{L}_i \text{s}^{-1}$ | *** $1.7 \pm 0.7$ | **** | $4.1 \pm 0.3$ |
| Sternohyoid |  |  |  |
| Muscle mass, g | 2.0 | ** 1.5 | 4.6 |
| Initial muscle length, mm | $21.2 \pm 0.07$ | $9.3 \pm 0.05$ | $15.3 \pm 0.01$ |
| Peak muscle strain, % | $2.5 \pm 0.7$ | $2.5 \pm 1.0$ | $2.7 \pm 0.4$ |
| Peak muscle velocity, $\text{L}_i \text{s}^{-1}$ | $2.7 \pm 0.8$ | $4.2 \pm 0.9$ | $4.8 \pm 0.4$ |
| Peak muscle mass-specific power, $\text{W kg}^{-1}$ | $79.1 \pm 25.6$ | * $223.6 \pm 87.2$ | $494.3 \pm 16.2$ |

\* Pressure, volume, and power values for Cb03 were measured for N = 7.
▲ Dorsal column muscle mass is included in the total epaxial muscle masses.
\*\* Bilateral muscle mass was estimated for Cb03 because individual was not available for muscle dissection.
\*\*\* Shortening of ventral (anal fin) muscle are reported here because hypaxial beads were not present in Cb01.
\*\*\*\* Hypaxial values were not recorded because hypaxial beads were not present in Cb03.

The dynamic endocast volume and buccal pressure measurements do introduce sources of error in the suction power estimates, as described in Camp et al. (2018). In brief, absolute volume measurements are overestimates, since they do not account for the presence of soft tissue or internal structures. However, the volume of these structures is consistent throughout the strike and should have little effect on the calculations for rate of volume change and subsequent power calculations. Rapid re-triangulation of dynamic endocast polygonal meshes may cause increased recorded rates of volume change, however, a low-pass, forward-backward Butterworth filter with a high cutoff frequency over the endocast buccal volume trace may produce underestimates of the actual rate of volume change. The intraoral pressure cannula only provides pressure readings at one location within the buccal cavity and does not capture variations in pressure during suction feeding (Muller et al., 1982; Van Wassenbergh, 2015). These estimates are likely underestimates of subambient pressure, since modeling of clariid catfishes and bluegill sunfish (Van Wassenbergh et al., 2015) and *in vivo* measurements (Tegge et al., 2020) suggested that highest subambient pressure occurred more caudally in the buccal cavity (Van Wassenbergh et al., 2005; Van Wassenbergh et al., 2006b). Additionally, our power calculations do not account for the forces required to overcome inertia or drag (Van Wassenbergh et al., 2015), yet studies of clariid catfishes and largemouth bass indicate that these forces are likely small compared to that required to overcome subambient pressure (Van Wassenbergh et al., 2005; Van Wassenbergh et al., 2015). Therefore, our values for instantaneous suction power are most likely to be underestimates.

### Determining peak gape

Suction feeding power, muscle shortening, and skeletal kinematics were all measured relative to the time of peak gape. Gape distance was measured as the distance between virtual locators on the rostralmost tips of the lower jaw and premaxilla. Peak gape is defined here as the maximum gape distance directly following the rapid increase in gape during the start of the strike. We calculated this by identifying the first frame at which there is a major change in the inflection of the gape distance curve from increasing to decreasing or in some cases minimal increasing. By taking the derivative of gape distance over time, we used 10% of the maximum rate of gape change as a threshold to isolate the first time point when the rate of gape change was below the threshold (Fig. S2). This method for determining peak gape differs from other suction power studies (Camp et al., 2015; Camp et al., 2018; Camp et al., 2020), but was chosen because of the high variability of gape distance traces in royal knifefish, e.g., gape curves with multiple peaks (double-strikes) or initial strikes followed by slow gradual gape expansion (slowly increasing plateau). Selecting the initial peak gape frame using the first instance of major inflection in gape distance yielded substantially better consistency of alignment of the expansion part of the gape cycle in this study (Fig. S2).

## RESULTS

In our study, royal knifefish were capable of generating very high suction power (Fig. 4). The neurocranium reached high magnitudes of elevation during the period of peak power (Fig. 5). Similarly, the epaxial muscle generated high strain and shortening velocity, reaching its shortest length during the period of peak power, and the sternohyoid shortened consistently across all strikes (Fig. 5). In addition, muscle mass-specific power was unusually high in the highest power strikes, reaching 535 W kg^-1^ in Cb03 and 800 W kg^-1^ in Cb04 (Fig. 4).

**Fig. 4.**
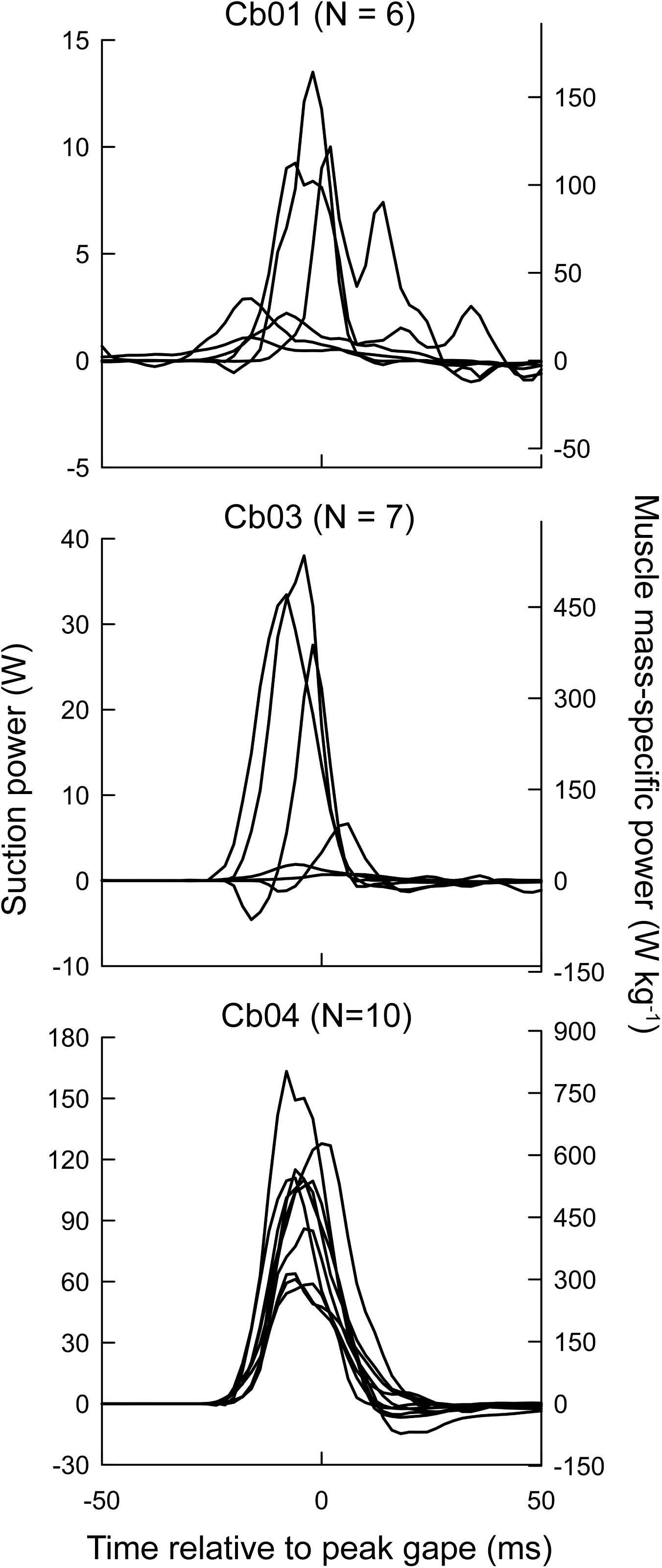
Suction power (W) and muscle mass-specific power (W kg^-1^) in three royal knifefish. Absolute suction power (left y-axis) is plotted relative to peak gape for each strike in each individual. The right y-axis shows muscle mass-specific power, which divides suction power by the combined mass of the epaxial and hypaxial shortening regions and the sternohyoid.

**Fig. 5.**
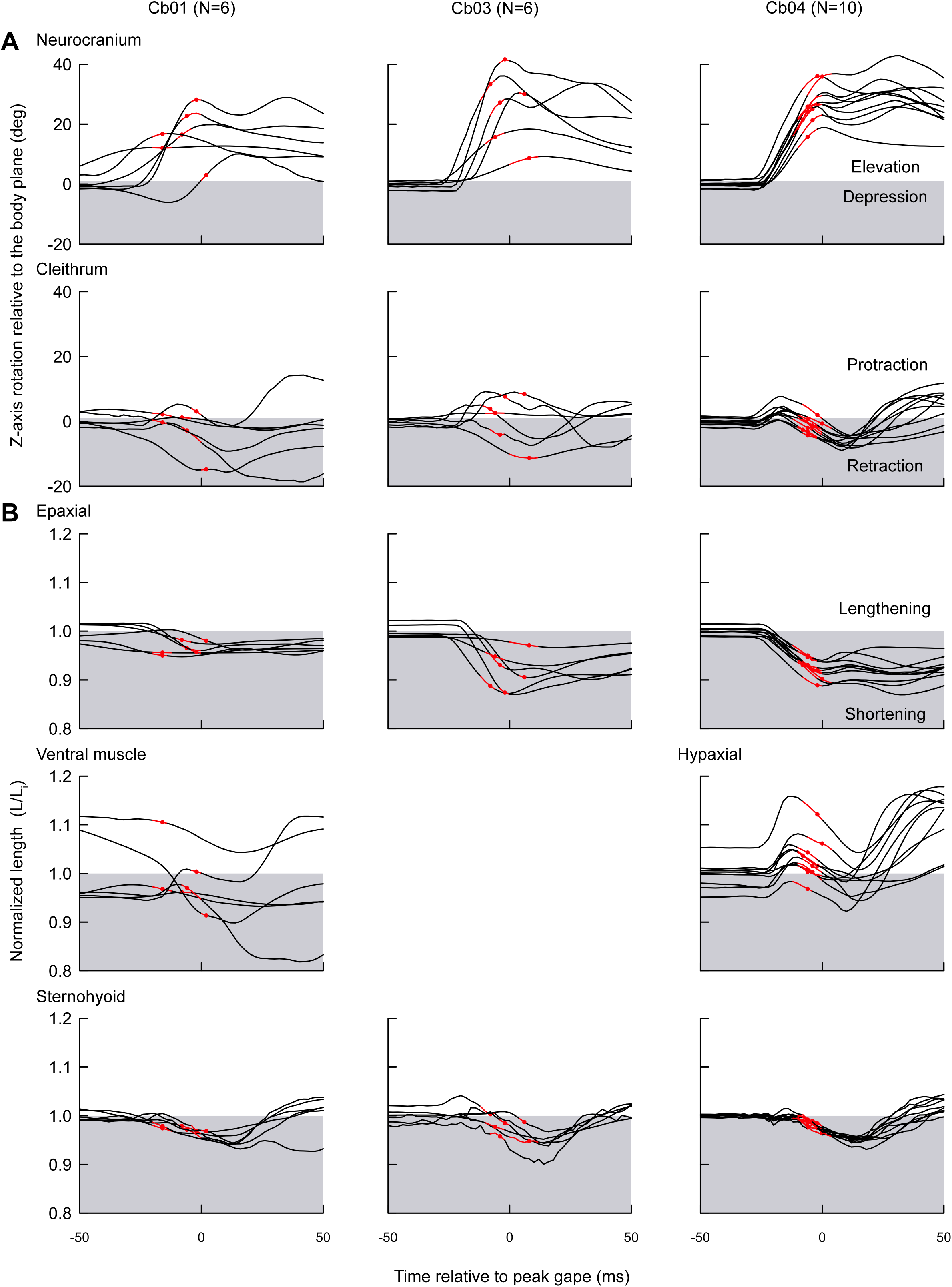
Neurocranium and cleithrum rotations relative to the body plane and muscle length changes during buccal expansion in royal knifefish. Data from each individual are shown separately, with the period of peak power (within 25% of maximum) highlighted in red and the time of peak power marked with a red dot. (A) Z-axis rotation (relative to initial values) of the neurocranium (row 1) and left cleithrum (row 2) relative to the body plane. Positive values represent elevation or protraction (white region), while negative values represent depression or retraction (shaded region). (B) For each muscle (bottom three rows), muscle length was normalized by its mean initial length (L_i_). Values below 1 (shaded region) represent shortened muscle, while those greater than 1 (white region) represent lengthened muscle relative to L_i_.

### Inter-individual variation

Cb04 produced substantially higher power strikes than Cb01 and Cb03 (Fig. 4). The mean peak power for Cb04 was 15 times greater than Cb01 and 6 times greater than Cb03 with the most powerful strikes reaching 13.5 W for Cb01, 38.0 W for Cb03, and 163.3 W for Cb04. The substantially larger suction power in Cb04 resulted from both greater subambient buccal pressure and faster rate of buccal volume change (Table 1). In Cb04, mean peak subambient pressure was approximately 5.6 times greater than that of Cb01 and nearly 3 times greater than that of Cb03, and mean peak rate of volume change was approximately 3 times greater than Cb01 and Cb03 (Table 1). Unlike Cb04, the difference in suction power between Cb01 and Cb03 was largely due to the difference in mean peak subambient buccal pressure, which was nearly two times larger in Cb03 compared to Cb01 (Table 1). Because of these differences among individuals, results are reported separately for each individual, with means and s.e.m. (Table 1).

### Skeletal kinematics

The neurocranium consistently elevated (rotated dorsally) relative to the body plane, across all strikes in all individuals (Fig. 5). During suction expansion, as the vertebral column extended from curved to straight, the cranialmost vertebrae elevated with the neurocranium (Fig. 2A,B). Due to placement of our body plane, the neurocranium JCS captures a combination of neurocranium elevation at the craniovertebral joint and vertebral column extension. While the magnitude of rotation was sensitive to location of the body plane, the neurocranium and anterior vertebral column elevated notably and consistently in all three individuals regardless of the body plane’s location. The mean maximum elevation measured during the period of peak power was 17.0 ± 3.6 deg for Cb01, 26.9 ± 4.9 deg for Cb03, and 27.9 ± 1.7 deg for Cb04. For some strikes, the neurocranium showed a pattern of initial elevation, slight depression, and then continued elevation at the end of pectoral girdle retraction. The initial phase of neurocranial elevation occurred during the period of peak power (shown in red in Fig. 5) and during epaxial muscle shortening (Fig. 5).

Cleithral retraction (caudoventral rotation) relative to the body plane was consistent within Cb04 strikes, but highly variable in Cb01 and Cb03 (Fig. 5). During Cb04 strikes, the cleithrum initially protracted (craniodorsal rotation), followed by the start of retraction prior to the period of peak power, and a steady, continued retraction through the period of peak power (Fig. 5). It should be noted that cleithral protraction occurs relative to the body plane; the cleithra are not protracting relative to the neurocranium but are instead being pulled dorsally by vertebral column extension, causing hypaxial lengthening. Maximum cleithrum retraction in Cb04 averaged -3.1 ± 0.6 deg during the period of peak power and increased to an average of -6.4 ± 0.5 deg after the period of peak power. Cb01 and Cb03 showed variability in timing and did not always retract during the period of peak power in their strikes. The magnitude of cleithral protraction and retraction was also highly variable in Cb01 and Cb03 (Fig. 5), with mean peak retractions of -3.0 ± 2.7 deg and 0.8 ± 3.0 deg, respectively, during the period of peak power.

### Muscle length changes and muscle power

The epaxial and sternohyoid muscles consistently shortened prior to and during peak power in all individuals (Fig. 5). However, the magnitude and pattern of epaxial shortening varied across individuals. Epaxial muscles shortened across all of the measured subregions in Cb03 and Cb04, and all but the caudalmost subregion of Cb01 (Fig. 6). Mean peak whole-muscle strain in the epaxials during the period of peak power was similar between Cb04 and Cb03 (8.0 ± 0.7 % L_i_ and 7.9 ± 1.4% L_i_, respectively), as was mean peak muscle shortening velocity during the period of peak power (4.7 ± 0.1 L_i_ s^-1^ and 4.6 ± 1.5 L_i_ s^-1^, respectively). Epaxial strains during the period of peak power were lower in Cb01, less than half that of Cb03 and Cb04 (Table 1). Mean peak epaxial strain during the period of peak power was lowest in the cranialmost region (below 5% strain in all individuals) and the highest at approximately one-half to three-fourths of the distance between the craniovertebral joint to the dorsal fin (3-6 cm, 4-7 cm, and 6-8 cm caudal of the craniovertebral joint in Cb01, Cb03, and Cb04, respectively) (Fig. 6; Fig. S1).

**Fig. 6.**
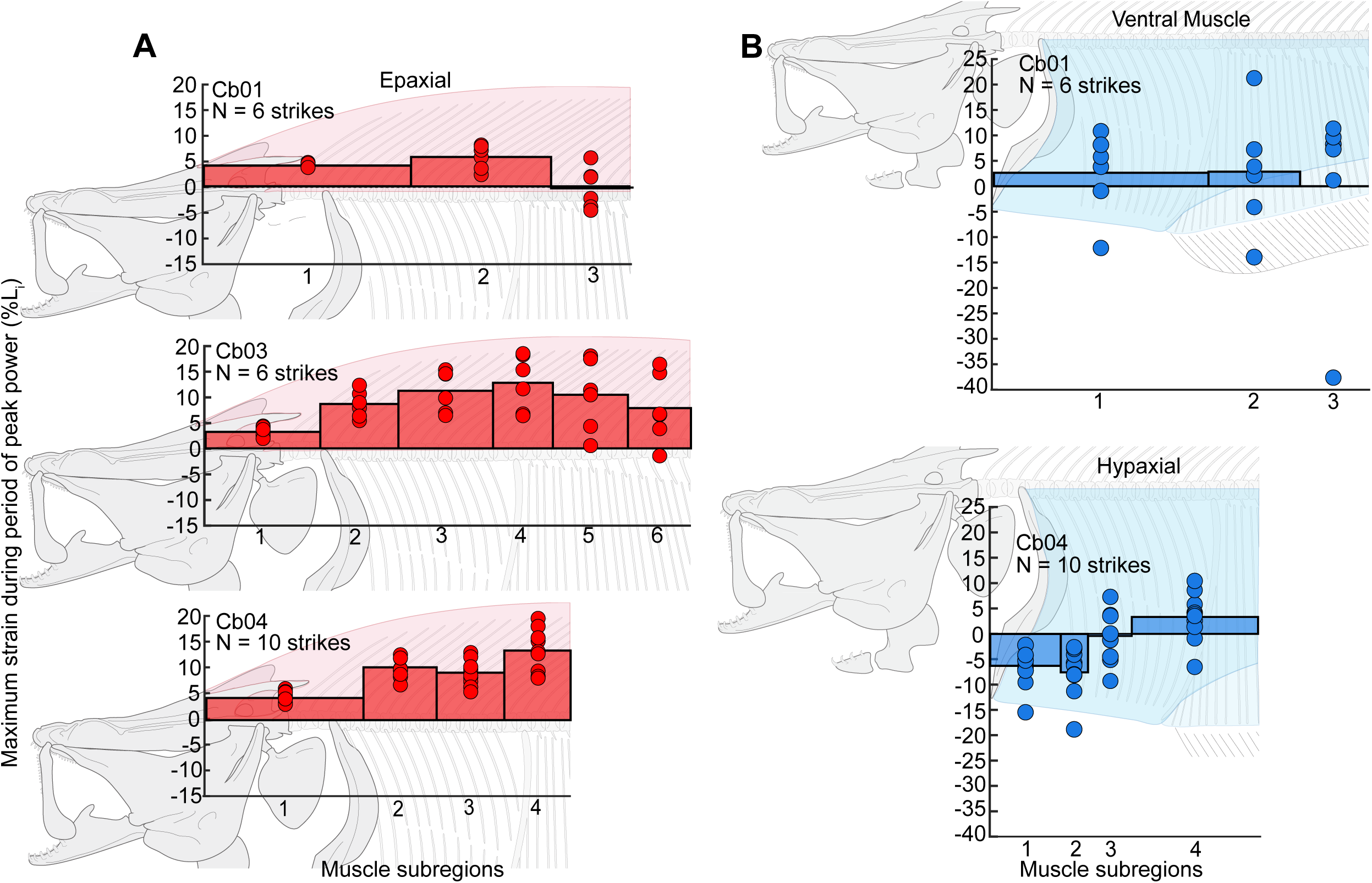
Maximum epaxial and hypaxial muscle strain along the body. (A) Epaxial strain and (B) ventral (anal fin) muscle and hypaxial strain during the period of peak power were calculated as percent change in length relative to the mean initial length (L_i_) in subregions along the cranial half of the body. Maximum strain for each strike (red or blue circles) and mean strain (red or blue bars) across all strikes are shown for each subregion with positive values representing muscle shortening. The width of the bar reflects the craniocaudal length of each muscle subregion. Note that Cb03 did not have ventral or hypaxial strain data.

Similar shortening behaviors were seen in the dorsal column of muscle of Cb04, the only individual with beads implanted in this muscle (Fig. S3). This column of muscle is dorsal to and separate from the epaxial. It inserts on the neurocranium and extends along the length of the body. The dorsal column of Cb04 shortened at the same time as the corresponding region of the epaxial muscle (three cranialmost subregions), reaching 5.8 ± 0.4 % L_i_ strain and muscle shortening velocity of 3.4 ± 0.1% L_i_ s^-1^ (Fig. S3). The caudalmost subregion showed the greatest magnitude of strain.

In the hypaxial musculature of Cb04, a consistent pattern of lengthening then shortening prior to the period of peak power occurred in all recorded strikes (Fig. 5). Early lengthening across the full extent of the marked hypaxial muscle was so great in Cb04 that, during peak power, it shortened with a mean peak velocity of 4.1 ± 0.1 L_i_ s^-1^, while the muscle length was still longer relative to its initial length (-1.1 ± 0.4% L_i_) (Table 1). The caudalmost subregion of the hypaxial muscle shortened during the period of peak power, while the cranialmost subregions lengthened (Fig. 6). Unlike Cb04, ventral muscle beads in Cb01 were implanted in the anal fin musculature, and no beads were implanted in the hypaxial muscle of Cb01 or Cb03 (Fig. S1). The shortening patterns and strain of the ventral muscle beads in Cb01, during the period of peak power, were highly variable (1.8 ± 2.8% L_i_) (Fig. 5, Table 1).

The sternohyoid shortened with a consistent pattern in all strikes and across individuals, with mean strains of 2.5-2.7% during the period of peak power (Table 1, Fig. 5). Sternohyoid shortening began prior to and continued through the period of peak power, with higher shortening velocities occurring during peak power. Magnitudes of strain in the sternohyoid were similar across individuals, but sternohyoid shortening velocity during the period of peak power was up to two times higher in Cb04 and Cb03 compared to Cb01 (Table 1).

Royal knifefish generated high muscle mass-specific power, which we calculated by dividing the maximum instantaneous suction power of each strike by the total mass of musculature shortening during peak power generation (epaxial, hypaxial and sternohyoid). For these muscles (0.2035 kg combined for Cb04) to produce the highest power strike recorded (163.3 W), they would have needed to generate 802.5 W kg^-1^ of power. The next three highest power strikes recorded for Cb04 are estimated to have required 628.0 W kg^-1^, 565.0 W kg^-1^, and 545.0 W kg^-1^. The maximum peak muscle power in Cb01 was much lower (164.2 W kg^-1^), but the muscle power in the highest power strike in Cb03 (535.3 W kg^-1^) was within the range of muscle power generated across all Cb04 trials (289.3 W kg^-1^ -802.5 W kg^-1^).

## DISCUSSION

The massive epaxial muscles of royal knifefish account for >25% of body mass and during suction feeding they shortened considerably and rapidly, generating large neurocranial elevation and vertebral extension. These results agree with our predictions based on body shape and postcranial morphology. During the most powerful strikes, the hypaxials also shortened and, together with the epaxials, generated over 95% of the power for suction expansion with muscle power output of up to 800 W kg^-1^. The magnitude of sternohyoid muscle shortening was consistent across all strikes, while the magnitude of axial muscle shortening was more variable during lower power strikes. This suggests that the sternohyoid muscle may contribute a greater proportion of power in lower power strikes. Likely driven by their large neurocranial elevation and rapid epaxial shortening, royal knifefish generated much higher rates of buccal expansion, subambient intraoral pressure, and suction power than those previously measured in other species (Table 2). For the purpose of this discussion, we will compare the highest performing individuals, using them as a proxy for the relative capabilities of each species (see *Variation in suction power* section).

**Table 2.**
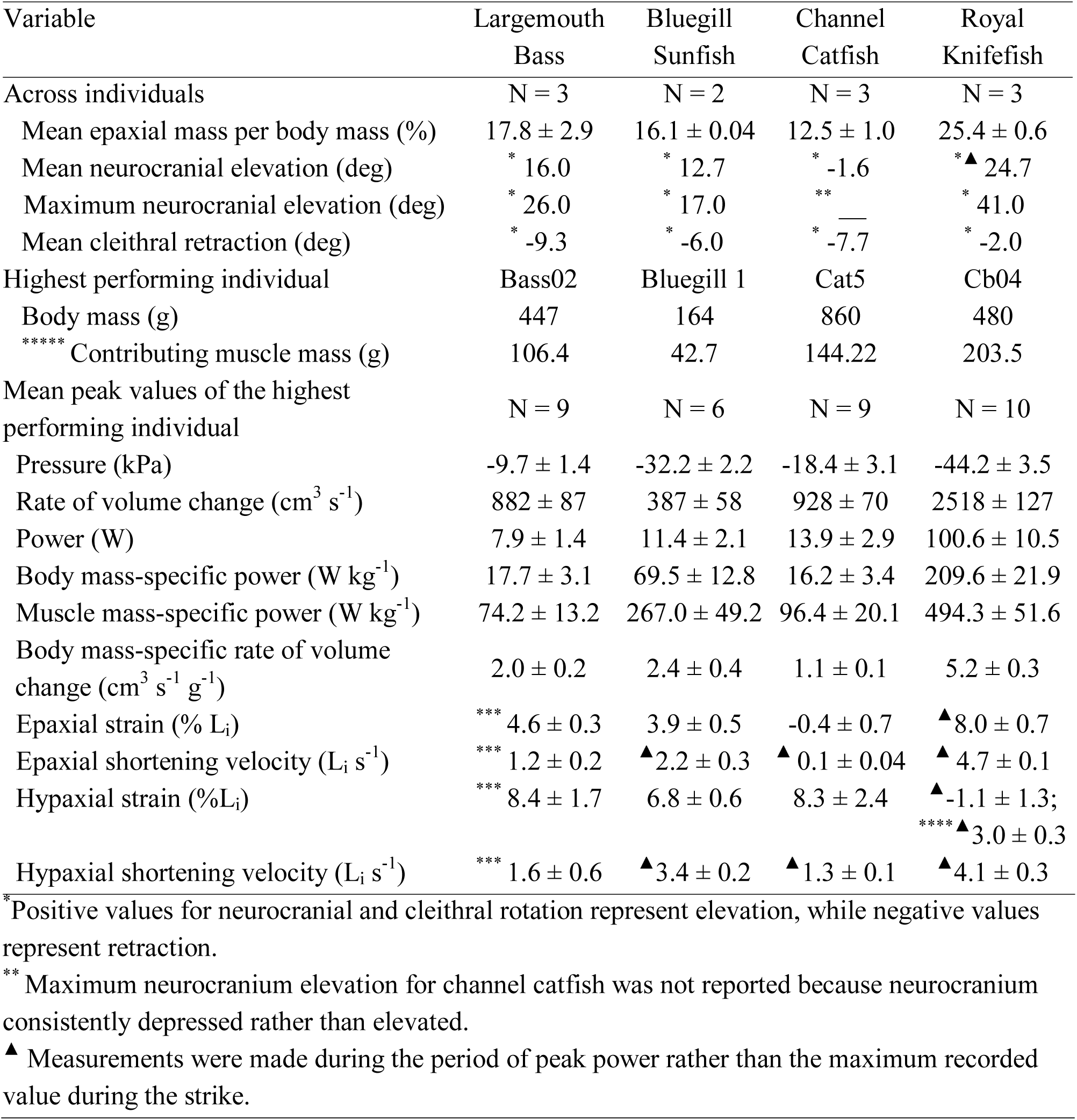
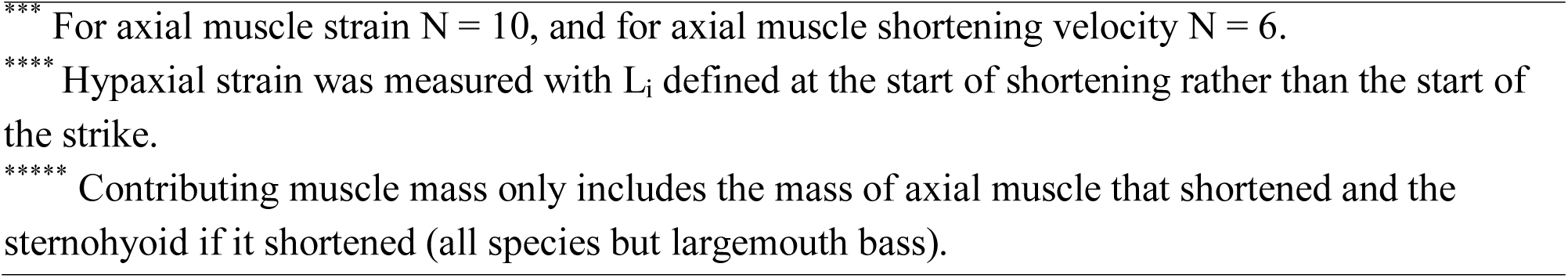
Comparative measurements for largemouth bass (Micropterus salmoides), bluegill sunfish (Lepomis macrochirus), channel catfish (Ictalurus punctatus), and royal knifefish (Chitala blanci). Data shown for royal knifefish are from this study, and data for other species are from previously published datasets (largemouth bass data from Camp et al., 2015; bluegill sunfish data from Camp et al., 2018; and channel catfish data from Camp et al., 2020). Where error values are included, they are the s.e.m (standard error of measurement).

| Variable | Largemouth Bass | Bluegill Sunfish | Channel Catfish | Royal Knifefish |
| --- | --- | --- | --- | --- |
| Across individuals | N = 3 | N = 2 | N = 3 | N = 3 |
| Mean epaxial mass per body mass (%) | 17.8 ± 2.9 | 16.1 ± 0.04 | 12.5 ± 1.0 | 25.4 ± 0.6 |
| Mean neurocranial elevation (deg) | * 16.0 | * 12.7 | * -1.6 | *<sup>▲</sup> 24.7 |
| Maximum neurocranial elevation (deg) | * 26.0 | * 17.0 | ** — | * 41.0 |
| Mean cleithral retraction (deg) | * -9.3 | * -6.0 | * -7.7 | * -2.0 |
| Highest performing individual | Bass02 | Bluegill 1 | Cat5 | Cb04 |
| Body mass (g) | 447 | 164 | 860 | 480 |
| ***** Contributing muscle mass (g) | 106.4 | 42.7 | 144.22 | 203.5 |
| Mean peak values of the highest performing individual | N = 9 | N = 6 | N = 9 | N = 10 |
| Pressure (kPa) | -9.7 ± 1.4 | -32.2 ± 2.2 | -18.4 ± 3.1 | -44.2 ± 3.5 |
| Rate of volume change (cm<sup>3</sup> s<sup>-1</sup>) | 882 ± 87 | 387 ± 58 | 928 ± 70 | 2518 ± 127 |
| Power (W) | 7.9 ± 1.4 | 11.4 ± 2.1 | 13.9 ± 2.9 | 100.6 ± 10.5 |
| Body mass-specific power (W kg<sup>-1</sup>) | 17.7 ± 3.1 | 69.5 ± 12.8 | 16.2 ± 3.4 | 209.6 ± 21.9 |
| Muscle mass-specific power (W kg<sup>-1</sup>) | 74.2 ± 13.2 | 267.0 ± 49.2 | 96.4 ± 20.1 | 494.3 ± 51.6 |
| Body mass-specific rate of volume change (cm<sup>3</sup> s<sup>-1</sup> g<sup>-1</sup>) | 2.0 ± 0.2 | 2.4 ± 0.4 | 1.1 ± 0.1 | 5.2 ± 0.3 |
| Epaxial strain (% L<sub>i</sub>) | *** 4.6 ± 0.3 | 3.9 ± 0.5 | -0.4 ± 0.7 | <sup>▲</sup> 8.0 ± 0.7 |
| Epaxial shortening velocity (L<sub>i</sub> s<sup>-1</sup>) | *** 1.2 ± 0.2 | <sup>▲</sup> 2.2 ± 0.3 | <sup>▲</sup> 0.1 ± 0.04 | <sup>▲</sup> 4.7 ± 0.1 |
| Hypaxial strain (%L<sub>i</sub>) | *** 8.4 ± 1.7 | 6.8 ± 0.6 | 8.3 ± 2.4 | <sup>▲</sup> -1.1 ± 1.3;<br>****<sup>▲</sup> 3.0 ± 0.3 |
| Hypaxial shortening velocity (L<sub>i</sub> s<sup>-1</sup>) | *** 1.6 ± 0.6 | <sup>▲</sup> 3.4 ± 0.2 | <sup>▲</sup> 1.3 ± 0.1 | <sup>▲</sup> 4.1 ± 0.3 |

\* Positive values for neurocranial and cleithral rotation represent elevation, while negative values represent retraction.
\*\* Maximum neurocranium elevation for channel catfish was not reported because neurocranium consistently depressed rather than elevated.
<sup>▲</sup> Measurements were made during the period of peak power rather than the maximum recorded value during the strike.
\*\*\* For axial muscle strain $N = 10$ , and for axial muscle shortening velocity $N = 6$ .
\*\*\*\* Hypaxial strain was measured with $L_i$ defined at the start of shortening rather than the start of the strike.
\*\*\*\*\* Contributing muscle mass only includes the mass of axial muscle that shortened and the sternohyoid if it shortened (all species but largemouth bass).

### Epaxial muscle shortening and neurocranial elevation

During suction feeding, royal knifefish substantially elevated their neurocranium and cranialmost vertebrae, fully straightening their vertebral column (Fig. 2). Mean maximum neurocranial elevation during the period of peak power in royal knifefish exceeded the mean maximum neurocranial elevation values previously measured in largemouth bass, bluegill sunfish, and channel catfish (Table 2). The highest values of neurocranial elevation during the period of peak power recorded in each royal knifefish individual (28-42 deg) were within the range observed in Commerson’s frogfish (*Antennarius commerson*), a genus known for its exceptionally large suction expansion (Camp, 2021; Longo et al., 2016). Our results are consistent with the predictions that a combination of the initially depressed neurocranium and ventrally flexed vertebral column increased the range of neurocranial motion used during suction feeding. Interestingly, these are anatomical traits shared by the frogfish (Camp, 2021) but not all species with extremely high cranial elevation (Lauder and Liem, 1981; Van Wassenbergh et al., 2008).

In royal knifefish, the epaxial musculature shortened along at least 60-70% of the body length, with high strain and shortening velocity. When comparing the highest performing individual of each species, the mean peak epaxial muscle strain during the period of peak power of royal knifefish was approximately two times the absolute peak epaxial strain (which occurred after the period of peak power) of largemouth bass and bluegill sunfish (Table 2). Similarly, the mean peak epaxial shortening velocity during the period of peak power was more than two times higher in the highest performing individual in royal knifefish than in largemouth bass and bluegill sunfish (Table 2). While the maximum shortening velocity (V_max_) for royal knifefish epaxials is unknown, V_max_ for largemouth bass has been measured as 11 L_i_ s^-1^ (Coughlin and Carroll, 2006), with 3-4 L_i_ s^-1^ as the range expected for optimal power output. Epaxial shortening velocity during the period of peak power in the highest performing royal knifefish was 4-5 L_i_ s^-1^, suggesting that the epaxials may be shortening at or near the range for optimal power output.

### Hypaxial muscle shortening and cleithral retraction

In Cb04, the cleithrum consistently protracted then retracted, while the hypaxial muscles lengthened then shortened. Cleithral protraction and hypaxial lengthening corresponded to the start of neurocranial elevation and vertebral column extension, which likely pulled rostro-dorsally on the cleithrum relative to the body plane, substantially lengthening the hypaxials. This pattern of cleithral protraction and hypaxial lengthening prior to cleithral retraction and hypaxial shortening has not been observed in ray-finned fishes previously studied with XROMM. Although this rostro-dorsal motion was measured as cleithral protraction, the cleithrum did not appear to protract relative to the neurocranium and we did not observe a reduction in buccal volume. Because the vertebral column remained partially extended after peak extension, shortening of the hypaxials back to just their initial length still retracted the cleithrum, on average, -6.4 ± 0.2 deg past its initial position (Fig. 5). Compared to the highest performing individual of each species, royal knifefish had greater mean peak hypaxial shortening velocity during the period of peak power than bluegill sunfish and more than 2.5 times that of largemouth bass and channel catfish (Table 2).

While the ventral muscle markers in Cb01 were implanted ventral to the hypaxial muscle, in the anal fin muscle (Fig. S1), the muscle length traces seemed to align with the cleithral retraction patterns as in Cb04 (Fig. 5). This suggests that anal fin data may still be reflective of hypaxial strain, but possibly more variable in lower power strikes.

### Relative contributions of the epaxial and hypaxial muscles

Our results suggest that the epaxial muscles are generating a greater portion of suction power than the hypaxial muscles in royal knifefish. First, mean epaxial muscle mass was 1.8 times greater than the hypaxial muscle mass, and so was capable of greater power output (Table 1). Second, in Cb01 and Cb03, cleithrum retraction—and presumably hypaxial shortening—were inconsistent, while neurocranial elevation and epaxial shortening were large and consistent (Fig. 5; Table 1). Although Cb04 used consistent cleithral retraction and hypaxial shortening, the magnitude and speed of hypaxial strain was less than half of epaxial strain (Table 1, Table 2). These data support the conclusion that the hypaxial muscles contributed less power, less consistently to suction expansion than the epaxial muscles.

### Sternohyoid muscle shortening and contributions to suction power

The timing and magnitude of sternohyoid shortening were relatively consistent across all strikes, irrespective of suction power and individual. All individuals had similar mean magnitudes of peak strain during the period of peak power, within 2.5-2.7 % L_i_ (Table 1). The sternohyoid shortened during the period of peak power and is electrically active during feeding strikes in congeneric species (Sanford and Lauder, 1989), which suggests that it actively contributed power to buccal cavity expansion. Consistent sternohyoid shortening similarly occurred during buccal cavity expansion in channel catfish, bluegill sunfish, striped surfperch (*Embiotoca lateralis*), and one clariid catfish (Camp et al., 2018; Camp et al., 2020; Lomax et al., 2020; Van Wassenbergh et al., 2007a). By contrast, in largemouth bass and several clariid catfishes, the sternohyoid did not shorten (or lengthen) during rapid suction expansion but rather acted as a stiff ligament that transmitted power from hypaxial musculature to produce hyoid depression (Camp and Brainerd, 2014; Van Wassenbergh et al., 2007a). Because the sternohyoid did not lengthen in royal knifefish, it also transmitted power generated from hypaxial shortening to facilitate hyoid depression and buccal expansion. This suggests that the sternohyoid has a dual function in royal knifefish, both transmitting power from the hypaxial muscle and generating power by shortening during the period of peak power.

The consistent pattern of sternohyoid shortening across all individuals suggests that the sternohyoid muscle may provide a greater proportion of muscle power in low performance strikes. To generate the suction power for the highest recorded strike in Cb04, we estimated that the musculature would need to generate 802.5 W kg^-1^. At this maximum muscle mass-specific power the sternohyoid in Cb04 (0.0046 kg) would be able to generate 3.7 W, which is within the range of the lowest power strikes recorded in Cb01 and Cb03. Similarly, the sternohyoid in Cb01 (0.002 kg) could produce up to 1.6 W of power, which is more than is necessary for suction expansion in the lowest power strike (1.1 W) from Cb01. It is still unlikely that the sternohyoid is the sole contributor since the neurocranium elevates and the epaxials shorten to some degree in all strikes (Fig. 5). Instead, the sternohyoid may make a greater contribution to generating suction power when epaxial shortening is low and hypaxial muscle shortening is inconsistent, as observed in Cb01 (Fig. 5). These results suggest that high power strikes depend nearly exclusively on axial muscle shortening, whereas a greater proportion of muscle power may come from the sternohyoid muscle in lower power strikes.

### Variation in suction power

Royal knifefish are capable of generating very high suction power, but we observed a wide range of power across the three individuals. Cb04 produced substantially higher power strikes, with mean peak suction power more than 15 times greater than Cb01 and 6 times greater than Cb03 (Fig. 4). The higher performance of Cb04 is partially explained by its body mass being more than twice the masses of Cb01 and Cb03 (Table 1), providing more muscle mass for power generation. When normalizing for body mass, there was substantial overlap in the mass-specific power in Cb03 and Cb04 (Fig. 7B), despite the non-overlapping ranges in absolute suction power (Fig. 7A). Additionally, it is possible that larger individuals generate more power per unit muscle mass if muscle power scales with positive allometry in royal knifefish as in other fish species (Carroll et al., 2009; Van Wassenbergh et al., 2007b). These features account for some of the variation between Cb03 and Cb04, suggesting that Cb04 may not merely be an exceptional individual.

**Fig. 7.**
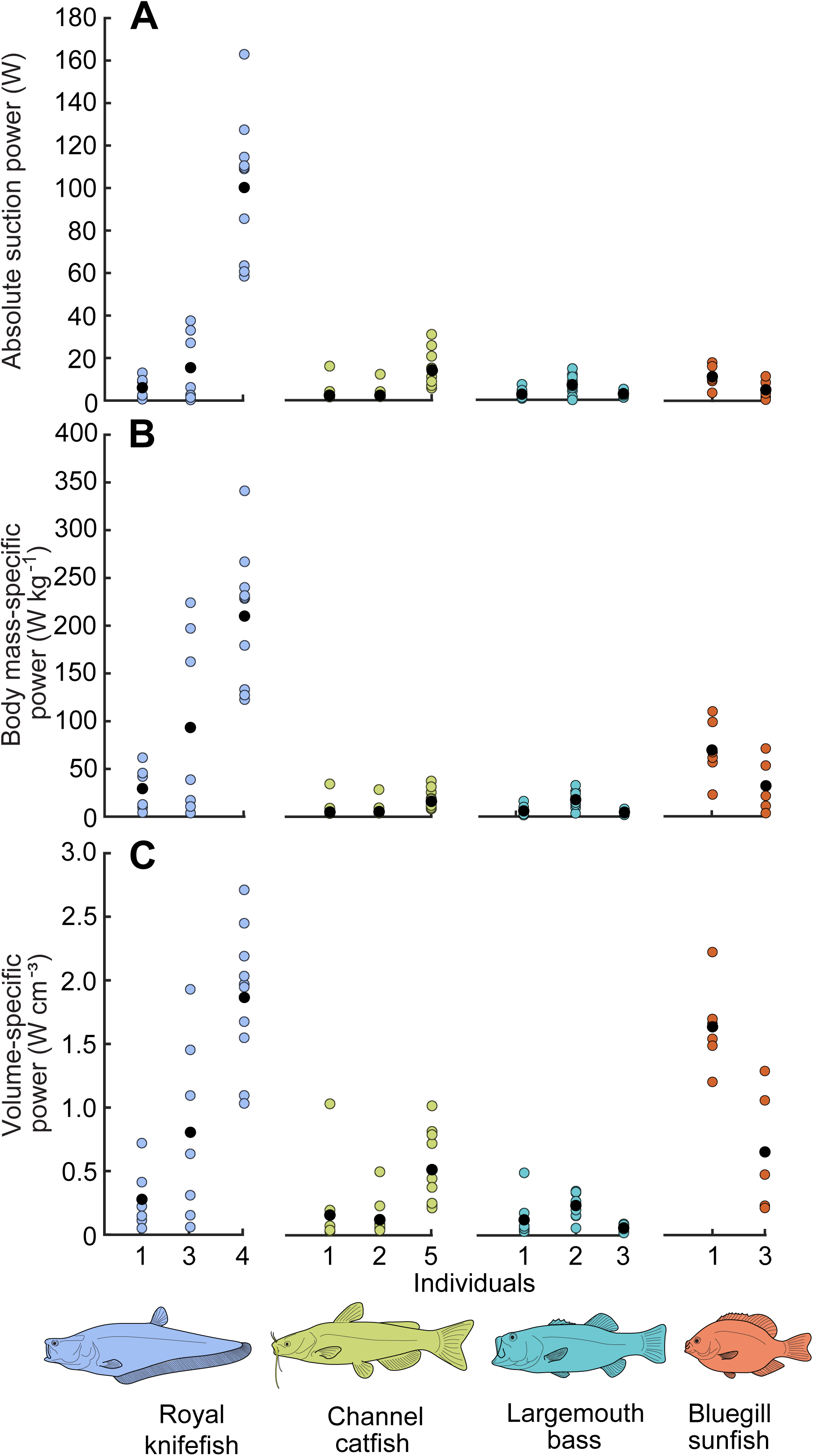
Comparison of suction power of royal knifefish to three other species. Data are shown for royal knifefish (n = 23 strikes, from this study), channel catfish (n = 24 strikes, data from Camp et al., 2020), largemouth bass (n = 29 strikes, data from Camp et al., 2015), and bluegill sunfish (n = 11 strikes, data from Camp et al., 2018). Power per strike (colored circles) and average power across all strikes from each individual (black circles) are shown. For all species, suction power was calculated as (A) the absolute magnitude of maximum suction power, (B) maximum suction power relative to the total body mass of the individual, and (C) maximum suction power relative to the maximum change in buccal volume for each strike.

However, body size does not completely explain intraspecific variation in power. While Cb03 had the smallest total body mass (78% of that of Cb01), it generated more than double the mean peak intraoral pressure and mean peak power compared to Cb01 (Fig. 4, Table 1). Interestingly, Cb01 and Cb03 had the same mean peak rate of buccal volume change. These results reflect that there is not a simple relationship between buccal volume change, intraoral pressure, and power, but instead a complex interaction between multiple factors, including gape size (morphologically and throughout the strike), initial buccal volume, magnitude of buccal volume change, timing of peak rate of buccal volume change, and timing of peak buccal cavity expansion (Van Wassenbergh et al., 2005; Van Wassenbergh et al., 2006a; Van Wassenbergh et al., 2006b). Motivation almost certainly contributed to this variation as well. Despite efforts to standardize prey type, prey size, and training, Cb04 responded better to training, was less timid when feeding in front of researchers, and was highly food motivated.

The variation among royal knifefish individuals is similar to what has been observed in bluegill sunfish, largemouth bass, and channel catfish. In all of these species, the highest performing individual generated mean body mass-specific suction power that was 2.0-2.6 times higher than the second highest performing individual (Camp et al., 2015; Camp et al., 2018; Camp et al., 2020). In addition, wide ranges of maximum suction power, intraoral pressure, and buccal volume change were observed, even when accounting for body or buccal volume size.

Both the present study and previous suction power studies are unlikely to have captured the maximum performance of any of these species, given the difficulty of eliciting maximum performance in lab-based studies with artificial environments and small sample sizes (Astley et al., 2013). Therefore, even the highest power strikes, such as those of Cb04, are conservative estimates of the suction power capacity of these species. Without having captured the true maxima of each species, conclusions from interspecies comparisons can only be drawn from the data that has been collected. Thus, while royal knifefish suction expansion appears impressively powerful compared to previously measured species, it is difficult to directly compare suction power capacity across species.

Within these limitations, comparing the data from the four species studied to date is a useful first step in exploring suction power across teleost fishes. When comparing across species, it may be most appropriate to compare high performing individuals to each other and lower performing individuals to each other. Within that context, all royal knifefish individuals outperformed largemouth bass, channel catfish, and bluegill sunfish individuals: Cb01 and Cb03 outperformed the lower performing individuals, just as Cb04 outperformed the highest performing individuals of those species (Fig. 7).

### Suction power, intraoral pressure, and buccal expansion

Royal knifefish generated higher suction power than the other species studied to date by producing both a greater magnitude of subambient intraoral pressure and a greater speed of buccal expansion (Camp and Brainerd, 2022). Of the four species, the mean peak intraoral pressure was greatest in royal knifefish, followed by bluegill sunfish, channel catfish, and largemouth bass (Table 2). Prior studies have not found consistent effects of body size on subambient buccal pressure, so we do not scale pressure here for body size (Carroll et al., 2004; Carroll et al., 2009). However, it is unclear how best to compare the rate of volume change among different sized individuals. If we simply compare the raw values across the highest performing individual within each species, the mean peak rate of buccal volume change was greatest in royal knifefish, more than 2.5 times that of largemouth bass and channel catfish, and more than 6.5 times that of bluegill sunfish (Table 2). If we normalize by body mass across the highest performing individual within each species, the mean peak body mass-specific rate of buccal volume change was still greater in royal knifefish, more than double that of bluegill sunfish and largemouth bass and almost five times that of channel catfish (Table 2). Thus, in comparison to the species previously studied with XROMM, the highest performing royal knifefish individual generated greater mean power by expanding its buccal cavity two times faster, relative to body mass, and generating at least 1.3 times greater buccal pressure magnitude.

Among the highest performing individuals of each species, royal knifefish generated a mean peak suction power approximately 8 times greater than channel catfish and bluegill sunfish and 13 times greater than largemouth bass (Fig. 7A; Table 2). When suction power was normalized by body mass or by maximum change in buccal volume—the difference between the volume of maximum buccal expansion and initial volume—then royal knifefish still outperformed the other three species but are more similar to bluegill sunfish (Fig. 7B,C).

### Muscle mass-specific power

For high performance strikes, royal knifefish depend on the axial musculature shortening at high velocities to produce large neurocranial elevation and rapid buccal expansion. At least 96.4% of the power for the highest power strike from Cb04 must have come from the axial musculature, based on the relative masses of the head and body muscles. If the major cranial muscles in Cb04 (sternohyoid, 4.6 g; levator arcus palatini, 1.64 g; levator operculi, 1.04 g; dilator operculi, 0.024 g) operated at the maximum muscle mass-specific power observed (802.5 W kg^-1^), the cranial muscles could generate 5.9 W of power. For Cb04, 5.9 W is just 3.6% of the maximum suction power and 4.6-5.3% of the next three highest power strikes. These results are consistent with the findings of previous studies, which have shown that cranial muscles are only capable of contributing a small proportion of the power necessary for high performance suction feeding and that the axial muscles are the primary source of suction power (reviewed in Camp and Brainerd, 2022).

Although we expected that royal knifefish would depend on their axial muscles to generate high power strikes, we did not expect the axial muscles to operate at such high muscle mass-specific power in the most powerful strikes. Compared to mean muscle mass-specific power outputs of the highest performing largemouth bass (74.2 ± 13.2 W kg^-1^), bluegill sunfish (267.0 ± 49.2 W kg^-1^), and channel catfish (96.4 ± 20.1 W kg^-1^), Cb04 achieved a greater mean muscle mass-specific power output of 494.3 ± 51.6 W kg^-1^ (Table 2), with a maximum of 802.5 W kg^-1^. This maximum muscle mass-specific power output is near or potentially beyond the expected limits for vertebrates (Altringham et al., 1993; Askew and Marsh, 2001; Curtin et al., 2005), suggesting several possible explanations: 1) we overestimated suction power; 2) we underestimated the longitudinal extent of the axial musculature that is contributing suction power; 3) there is power amplification, in which the muscle shortens before the skeletal elements begin to move, thereby loading serial elastic elements that release their energy while the muscle continues to contract as the bones move (Astley and Roberts, 2012).

In considering this first explanation, our suction power estimates are conservative because we measured buccal pressure in one rostral location and hydrodynamic modeling has shown that pressure can be even more subambient in the caudal buccal cavity (Van Wassenbergh, 2015). For the second, we dissected and included the mass of nearly 75% of the total epaxial and hypaxial length (Fig. S1), extending more caudally than our implanted marker set in order to provide a generous muscle mass estimate. The third possibility is power amplification, as is seen in the epaxial musculature of pipefishes and seahorses (Van Wassenbergh et al., 2008; Van Wassenbergh et al., 2014). This is an exciting potential explanation, but our results do not support this hypothesis. If power amplification were to create a catapult-like mechanism, we would expect to see gradual muscle shortening prior to the beginning of the strike to store elastic energy in the muscle and connective tissues (Astley and Roberts, 2012; Van Wassenbergh et al., 2008). However, there was no indication of muscle shortening prior to the beginning of neurocranial elevation nor pectoral girdle retraction (Fig. 5). We conclude that power amplification is unlikely the cause of such high muscle mass-specific muscle power estimates and that Cb04 was able to power suction feeding directly with 550-800 W kg^-1^ of muscle power in its four most powerful suction strikes.

### Concluding remarks

Compared to the three species previously studied with XROMM, royal knifefish are distinct in the morphology of their postcranial musculoskeletal system and their reliance on epaxial muscles for suction feeding. In royal knifefish, their epaxial muscles were greater in mass relative to the hypaxial muscles and shortened rapidly, producing a majority of suction power with rapid neurocranial elevation. We expect that species with similar morphology (including a ventrally flexed vertebral column, dorsoventrally deep epaxial muscles, few bones immediately caudal to the neurocranium, and a high proportion of epaxial muscle mass) can also produce high power strikes that are generated predominantly by epaxial muscle power and that utilize large neurocranial elevation. Our results support the growing evidence that postcranial morphology is important for understanding suction feeding mechanics, and that these feeding functions have likely shaped the evolution of the axial muscles and skeleton. Royal knifefish used nearly their entire body musculature to generate their most powerful strikes, broadening the morphological and phylogenetic range of suction feeding fishes known to power feeding with body muscles. However, the sternohyoid muscle likely contributed a greater proportion of power in the lowest performance strikes, demonstrating that the roles of cranial and axial muscles may vary not only across species, but also among feeding behaviors. Further studies examining the cranial and axial musculoskeletal systems—and their interaction—are needed to understand how the morphology of the whole body shapes the evolution and mechanics of suction feeding.

## ACKNOWLEDGEMENTS

We are grateful to Erika Tavares for research administrative support and manuscript edits, to Peter Falkingham and Stephen Gatesy for their assistance in developing the methods for the dynamic digital endocast, to David Baier for the XROMM Maya Tools, and to Jake Parsons, Noah Sims, and Paola Vazquez for their assistance tracking videos.

## DATA AVAILABILITY

X-ray video, pressure, and CT data and their essential metadata for this publication have been deposited in the XMAPortal (xmaportal.org), in the study “Knifefish Suction Feeding,” with the permanent identifier BROWN65. The data will be publicly available under CCBY 4.0 upon publication in the Public Data Collection “Royal Knifefish data for Li, Kaczmarek et al., 2022” with the full URL: https://xmaportal.org/webportal/larequest.php?request=CollectionView&StudyID=65&instit=BROWN&collectionID=22

