## Supplemental Figures for "Royal knifefish generate powerful suction feeding through large neurocranial elevation and high epaxial muscle power"

Supplemental Information

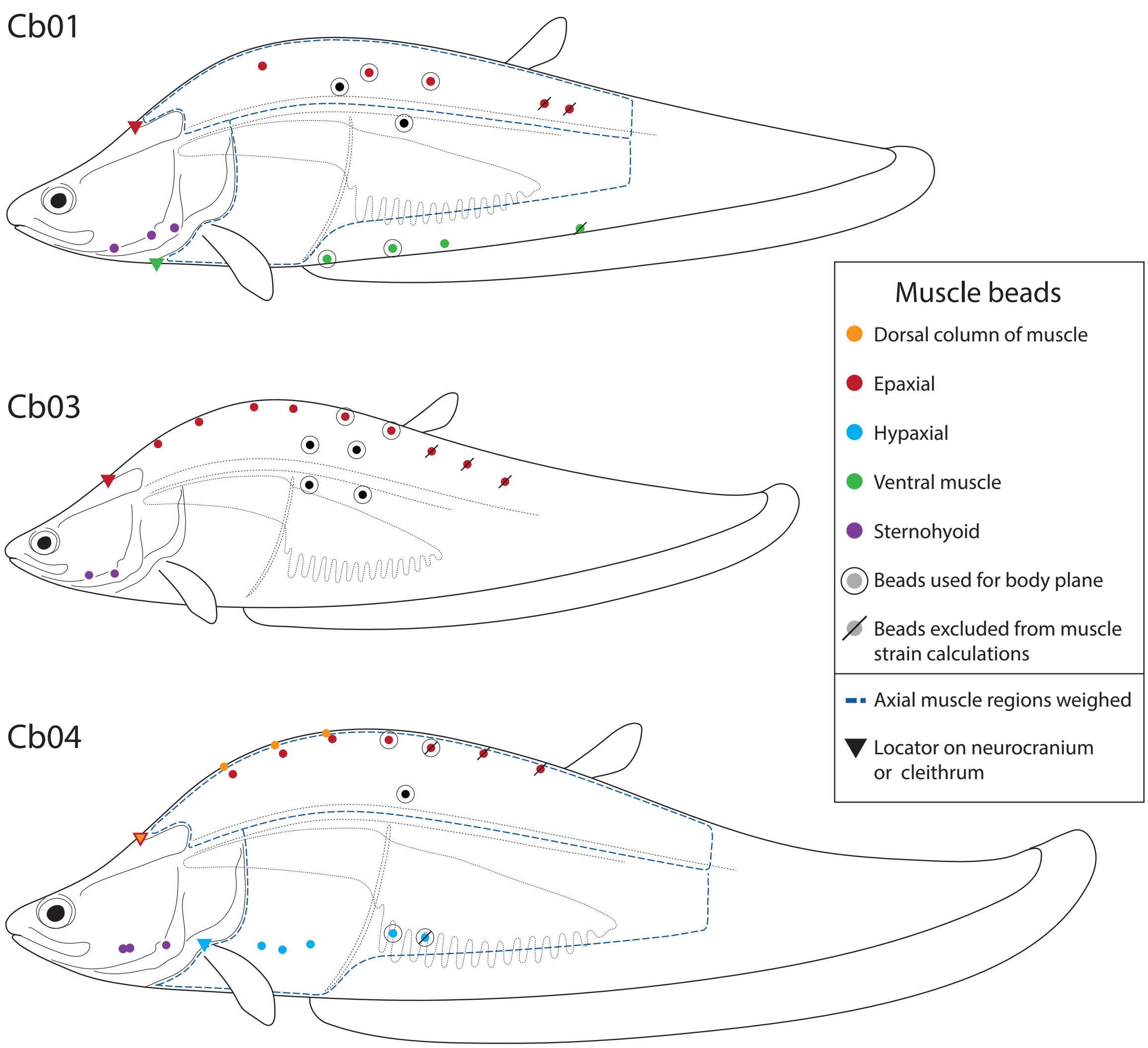


**Fig. S1. Intramuscular bead set for each individual.** Lateral whole-body illustrations are drawn proportional to the size of each individual. Intramuscular bead locations for the dorsal column (orange), epaxial (red), hypaxial (blue), ventral (anal fin) muscle (green), and sternohyoid (purple) are indicated with filled circles. Virtual locators (indicated with triangles) placed on the neurocranium and cleithrum were used to calculate muscle strain in the cranialmost subregions of the dorsal column, epaxial, hypaxial, and ventral (anal fin) muscles. The beads (in the epaxial, hypaxial, and ventral muscle) that were not included in the muscle length plots and muscle strain calculations have slashes through them (these caudal beads were not visible in the majority of strikes). Beads used to animate the body plane are highlighted with black circles. Dark blue dashed lines indicate the regions of epaxial and hypaxial muscles that were weighed. Note that Cb03 was not available for muscle dissection, so its axial muscle masses were estimated based on total body mass.

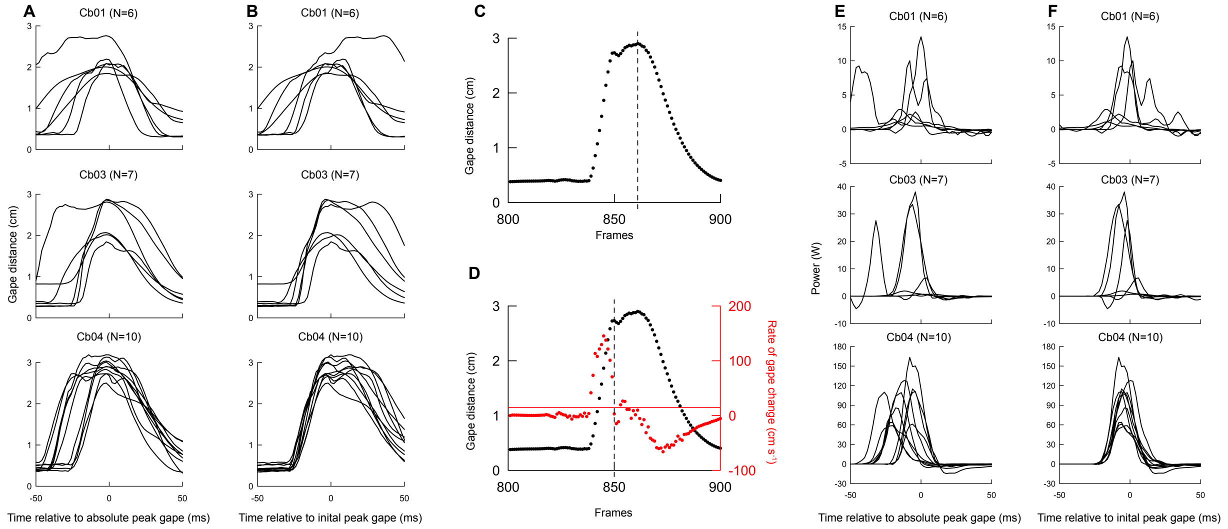


**Fig. S2. Comparison of measuring time relative to absolute peak gape and to initial peak gape.** (A,B) Gape distance for all strikes with time plotted relative to the timing of absolute peak gape and to the timing of initial peak gape, respectively. (C) The time of absolute peak gape (dashed vertical line) is the time of maximum gape distance in a trial. (D) The time of initial peak gape (dashed vertical line) is the first time point when the rate of peak gape change (red) is below 10% of the maximum rate of gape change (solid, horizontal red line). (E,F) Suction power for all strikes with time plotted relative to the timing of absolute peak gape and to the timing of initial peak gape, respectively. In this study, peak gape is defined as initial peak gape, not absolute peak gape.

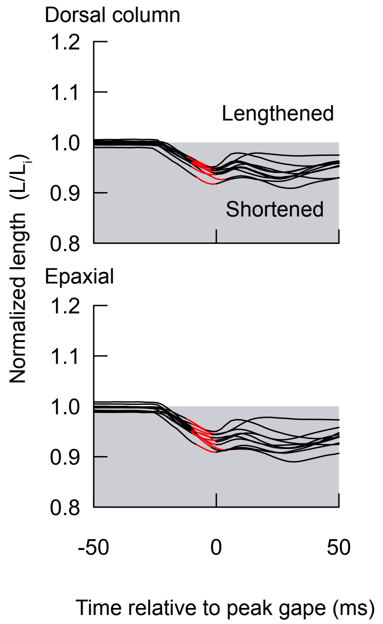


**Fig. S3. Muscle length changes in the dorsal column and epaxial muscle of Cb04.** Muscle length (black) normalized by the mean initial length ($L_i$) is plotted for the dorsal column (top) and epaxial muscles (bottom) for each Cb04 strike. Epaxial and dorsal column length changes are measured up to the third marker in each muscle, spanning the same extent of the body. Note that in Fig. 5, epaxial length change is measured up to the fourth marker. Values below 1 (shaded region) indicate that the muscle has shortened and values greater than 1 (white region) indicate that the muscle has lengthened relative to its initial length ($L_i$). The period of peak suction power (within 25% of maximum power) is highlighted in red for each strike.

**Video S1. Video of Cb04 feeding on a goldfish in the tunnel extension of a tank.** The video was recorded at 500 frames $s^{-1}$ and slowed down 16.67 times. A barrier is lifted, revealing a goldfish at the end of the tunnel. Cb04 approaches, slows down, and then strikes.
